# ER tethering and active transport govern condensate diffusion during hyperosmotic stress

**DOI:** 10.1101/2025.06.13.659610

**Authors:** Bisal Halder, Guoming Gao, Armin Ahnoud, Shelby Stakenas, Emily R. Sumrall, Nils G. Walter

## Abstract

**Background:** Hyperosmotic shock and the resulting cell volume compression are commonly experienced by organs such as the kidneys, causing rapid formation of hyperosmotic phase separation (HOPS) condensates in the cytoplasm and nucleoplasm. Although the tight relationship between hyperosmotic shock and condensation has been characterized, the dynamics of biomolecular condensates in hyperosmotically compressed cells and their regulatory mechanisms remain largely unknown.

**Results:** We used live-cell single-particle tracking (SPT) across different time scales to systematically characterize the dynamics of HOPS condensates formed by model protein mRNA decapping enzyme 1A (DCP1A). We found that HOPS condensates predominantly exhibited sub-diffusion rather than free diffusion, whereas some (∼2%) exhibited short super-diffusion. Using tools measuring spatial accessibility inside cells and fluorescence labels for specific cellular organelles, we further revealed the origins of sub-diffusion and super-diffusion as endoplasmic reticulum (ER) attachment and coupling to microtubule-dependent active transport, respectively. Further, we reconstructed an accessibility map of the hyperosmotically compressed cell from trajectories of genetically encoded multimeric nanoparticles (GEMs), revealing that the cytoplasm of a compressed cell remains highly accessible without significant local corrals.

**Conclusions:** In contrast to prior portrayals of the cytosolic space as static and constrained, our data suggest that the cytosol of a hyperosmotically compressed cell remains dynamic and accessible. Meanwhile, hyperosmotic and potentially other condensates can be spatially organized through docking to membrane structures, with intermittent episodes of long-range transport. These insights broaden our understanding of the physical environment within cells under hyperosmotic shock and provide a model for spatiotemporal organization of condensates via docking or coupling to existing cellular structures and processes.

## Background

Biomolecular condensates are membraneless, dynamic compartments that orchestrate the spatial and temporal organization of cellular biochemistry, playing essential roles in processes such as transcription, RNA metabolism, and signal transduction [1–3]. These structures typically arise via liquid–liquid phase separation (LLPS), a process governed by multivalent interactions among proteins and RNAs, and are highly sensitive to environmental cues including pH, temperature, and ionic strength. While in vitro studies have elucidated the molecular grammar underlying condensate assembly [4, 5], our understanding of how condensates form, function, and are regulated within living cells, especially under physiological stress, remains incomplete.

Hyperosmotic shock and the resulting cell volume compression are commonly experienced by organs such as the kidneys, which routinely encounter dramatic fluctuations in extracellular osmolarity due to their central role in water and electrolyte homeostasis [6]. In the renal medulla, cells are exposed to steep osmolarity gradients ranging from 300 mOsm/L in the cortex to over 1,200 mOsm/L in the inner medulla [7], creating conditions where osmotic fluctuations are essential for normal kidney function. The urine-concentrating mechanism generates these osmolarity fluctuations to enable cells to conserve or eliminate water and electrolytes, making osmotic stress adaptation particularly critical in cells located in the loop or the later section of the collecting duct, given their higher fluctuations in osmotic pressure [6].

This physiological osmotic stress causes rapid formation of hyperosmotic phase separation (HOPS) condensates in both the cytoplasm and nucleoplasm within seconds of osmotic compression [6, 8, 9]. HOPS represents a widespread cellular mechanism that affects a substantial fraction (∼10%) of the human proteome, with each phase-separating protein reversibly forming 300 or more condensates within as little as 10 seconds [9]. Importantly, nuclear HOPS condensates have profound implications for gene expression regulation [6], making this phenomenon particularly relevant to genomics research. For instance, HOPS-induced YAP condensates reorganize chromatin topology and recruit transcription machinery to activate specific gene expression programs [10]. Similarly, HOPS-induced CPSF6 condensates sequester cleavage and polyadenylation factors away from transcription termination sites [9], leading to widespread transcriptional readthrough and the production of stress-induced downstream-of-gene (DoG) transcripts from as many as 10% of all protein-coding genes in humans [11, 12]. These nuclear condensates thus establish genome-wide transcriptional responses that are fundamental to cellular adaptation and stress response mechanisms.

Although the relationship between hyperosmotic shock and condensation has been well characterized, the dynamics of biomolecular condensates in hyperosmotically compressed cells remain largely unexplored. Previous studies investigating condensate behavior have employed imaging durations of only a few seconds [6, 9], severely constraining observable dynamics and missing longer timescale behaviors and rare events. This methodological limitation has prevented the field from capturing the full spectrum of condensate dynamics, particularly rare but potentially important behaviors that occur beyond typical observation windows.

These dynamics are of critical importance since the physical environment within cells is widely believed to change dramatically upon hyperosmotic cell volume compression [13, 14], yet direct experimental evidence characterizing how condensates navigate this altered landscape is lacking. The prevailing assumption has been that cell compression leads to a crowded, static, and potentially corralled cytosolic environment that would restrict condensate mobility, but this hypothesis has not been rigorously tested through systematic analysis of condensate movement patterns.

Understanding condensate dynamics is critical because numerous important membraneless organelles, particularly in the nucleus, regulate gene expression and establish transcriptional programs throughout the genome [15]. Nuclear condensates such as nucleolus, transcriptional hubs, speckles, paraspeckles, histone locus body, PML body, and so on are central to genomics studies focused on understanding spatiotemporal control of gene expression [16–18]. The physical principles governing condensate behavior under osmotic stress likely extend beyond HOPS to influence the broader landscape of membraneless organelles that orchestrate genomic function, with implications for nuclear organization, transcriptional control, and cellular adaptation mechanisms [8].

In this study, we systematically investigate the dynamic behavior of HOPS condensates under hyperosmotic stress using live-cell single-particle tracking across extended timescales. By focusing on DCP1A as a model system, we reveal that HOPS condensates exhibit predominantly sub-diffusive motion through specific interactions with the endoplasmic reticulum, while a small fraction undergoes microtubule-dependent super-diffusive transport. Contrary to prevailing assumptions of a static, corralled cytosolic environment, our findings demonstrate that the cytosolic space of hyperosmotically compressed cells remains dynamic and accessible, with condensate organization achieved through docking to membrane structures rather than physical entrapment.

## Results

### HOPS condensates are significantly confined with occasional super diffusion

Previous studies of condensate dynamics under hyperosmotic shock have been constrained by brief imaging durations of only a few seconds [9], severely limiting our ability to capture the full spectrum of condensate behaviors and potentially missing rare but biologically significant events. When diffusion metrics are extracted using mean squared displacement analysis, distinct diffusion states—sub-diffusion, normal diffusion, and super-diffusion—exhibit significant overlap at early time lags [19, 20], requiring extended observation periods to achieve clear separation between these modes (Fig. 1a). To overcome this fundamental limitation and reveal the complete dynamic landscape of HOPS condensates, we implemented prolonged imaging protocols spanning approximately 6.5 minutes at 0.5 Hz, enabling systematic characterization of both common and rare diffusion behaviors. This extended temporal resolution proved essential for uncovering the complex dynamics that govern condensate behavior under hyperosmotic stress, providing the foundation for mechanistic insights that would otherwise remain hidden.

**Fig. 1.**
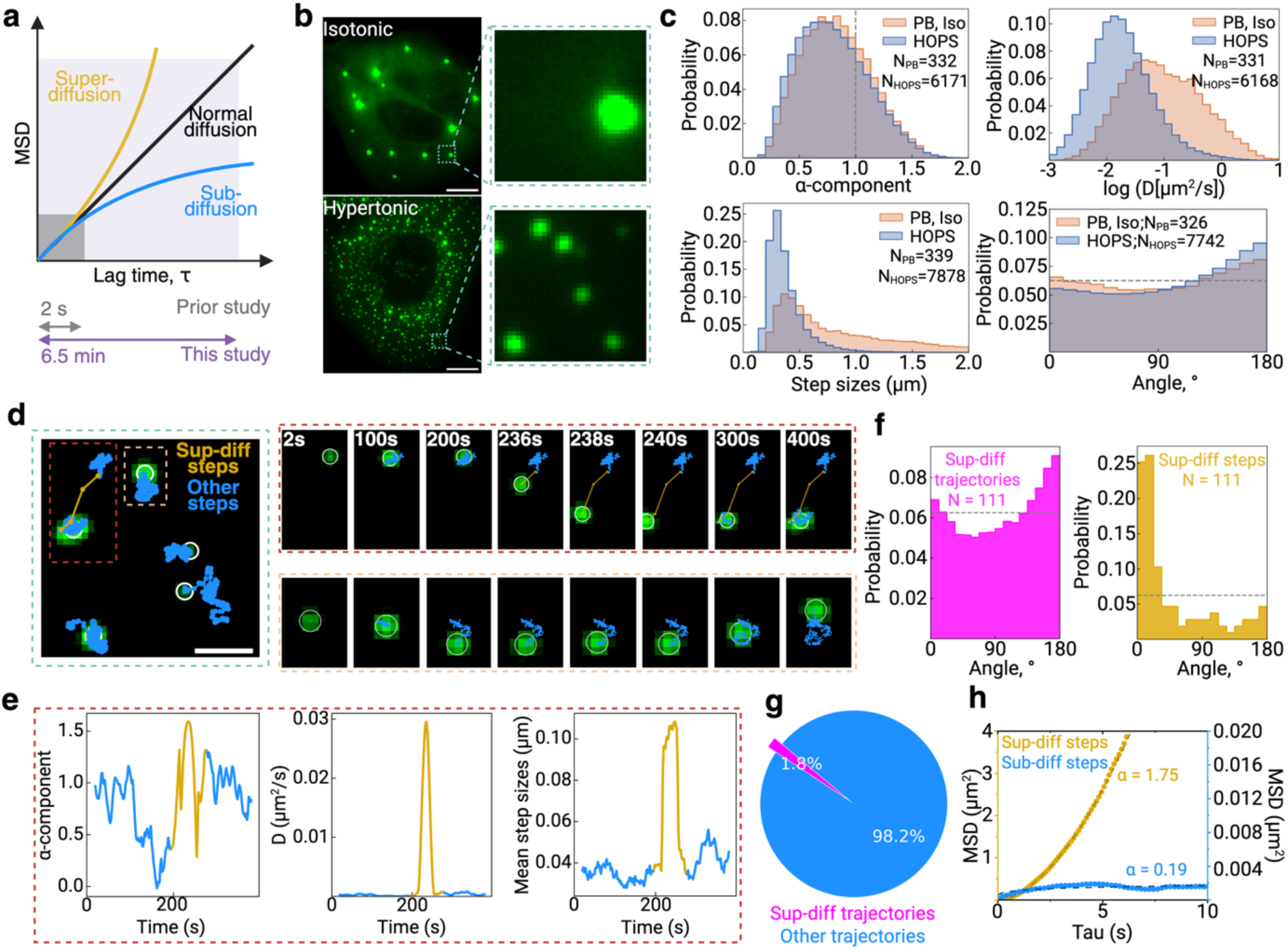
Hyperosmotic phase separation (HOPS) condensates undergo predominantly confined diffusion with occasional super diffusion. **a |** The mean squared displacement (MSD) as a function of lag time (τ) demonstrates the ability of extended time scales to differentiate between various diffusion modes. **b |** Representative images of U2OS cells expressing eGFP-labeled DCP1A under physiologically isotonic conditions (top) and hyperosmotic stress conditions (bottom). Enlarged views emphasize processing bodies (PBs) in isotonic conditions and hyperosmotic stress-induced HOPS condensates. Scale bar is 10 µm. **c |** Distributions of single-particle tracking parameters for PBs and HOPS condensates using a running-window analysis: top left, anomalous diffusion exponent (α); top right, diffusion coefXicient (D); bottom left, step size; and bottom right, angle between successive steps. **d |** A reconstructed high-resolution zoom-in of a representative region displays individual HOPS condensate trajectories exhibiting super diffusion (gold) and sub diffusion (blue), with time-lapse images of representative trajectories for each type of diffusion. Scale bar is 1 µm. **e |** Running-window analysis of a representative HOPS condensate trajectory in **d**, showing the change of α, D, and step size over time. **f |** Distribution of angles between adjacent steps within a HOPS condensate trajectory that contains a super diffusion session. **g |** Pie chart showing the relative proportions of HOPS condensate trajectories that contains a super diffusion session versus not. **h |** Comparison of MSD-τ curves between the sub-diffusive portions and the super diffusion portions of the same set of HOPS condensate trajectories.

Under hyperosmotic stress conditions (300 mM Na⁺), DCP1A undergoes a dramatic redistribution from 10-30 processing bodies per cell (200-800 nm radius) to 200-300 HOPS condensates per cell (150-350 nm radius) (Fig. 1b). Quantitative analysis using running window approaches revealed that HOPS condensates are significantly more confined than processing bodies under isotonic conditions, with the distribution of anomalous diffusion exponents (α) shifted toward substantially lower values (mean α = 0.73 for HOPS versus α = 0.8 for PBs). This increased confinement was further supported by diffusion coefficient analysis, which showed HOPS condensates exhibiting a distinct, narrow peak at logD = −1.81 µm^2^/s, compared to the broader distribution of PBs centered at logD = −1.27 µm^2^/s (Fig. 1c). Step size distributions provided additional compelling evidence, with HOPS condensates displaying a sharp, restricted peak at 0.33 μm versus the broader PB distribution centered at 0.40 μm. Most strikingly, angular analysis between consecutive steps revealed that HOPS condensates exhibit markedly more frequent directional reversals, with their distribution skewed toward 180°, indicating sub-diffusive, back-and-forth movement characteristic of trapped particles. These convergent lines of evidence establish that HOPS condensates exhibit a fundamentally more sub-diffusive dynamics than processing bodies, suggesting distinct underlying mechanisms governing their intracellular motions.

Despite their predominantly sub-diffusive nature, extended imaging unveiled a remarkable and previously undetected phenomenon: HOPS condensates undergo transient episodes of super-diffusive motion, characterized by rapid, directional “jumping” between distinct confined states (Fig. 1d). These super-diffusive events were captured through our running window analysis as sudden spikes in the anomalous diffusion component (α > 1), diffusion coefficient, and step size, occurring against the background of otherwise sub-diffusive motion (Fig. 1e). Angular distribution analysis of these super-diffusive trajectories revealed a distinct behavioral signature: while the overall population showed frequent directional reversals (sub-diffusion), super-diffusive steps exhibited a highly pronounced peak near 0°, indicating strong directional persistence during active transport events (Fig. 1f). Moreover, further quantification of these super diffusion event shows that only ∼2% of the HOPS condensate trajectory steps are on super diffusion throughout the whole 6.5 min observation window (Fig. 1g). Once in the “jumping” period, HOPS condensates can exhibit an ensemble α as high as 1.75, significantly higher than an ensemble α of 0.19 when outside the “jumping” period (Fig. 1h). This again validates that these “jumping” events are super diffusion.

Taken together, our extended imaging approach reveals that HOPS condensates exhibit a previously uncharacterized dual-mode dynamic behavior under hyperosmotic stress. The vast majority of condensates display significantly more confined, sub-diffusive motion compared to processing bodies under normal conditions, as evidenced by reduced anomalous diffusion exponents, restricted step sizes, and frequent directional reversals. In stark contrast, approximately 2% undergo rare but highly directional super-diffusive “jumping” events that facilitate transitions between spatially separated confined regions. This discovery of regulated mobility, where predominant confinement is punctuated by occasional bursts of active transport, fundamentally challenges the prevailing view of condensates as static entities under osmotic compression and establishes a new paradigm for understanding how these stress-induced assemblies navigate the intracellular environment. Importantly, these findings are robust across imaging frequencies, as identical results were obtained using both 0.5 Hz and 50 Hz acquisition rates (Supplementary Fig. 1), confirming that extended duration rather than high temporal resolution is the critical factor for capturing the full spectrum of condensate dynamics.

### Super diffusion of HOPS condensates is microtubule-dependent

Super-diffusion represents an energy-dependent process typically mediated by motor proteins that convert ATP hydrolysis into mechanical force for active cargo transport [21, 22], suggesting that the super-diffusive behavior observed in DCP1A HOPS condensates reflects motor-driven transport rather than passive diffusion. To test this hypothesis and identify the specific cytoskeletal pathway underlying super-diffusion, we employed targeted pharmacological disruption of the two major cytoskeletal networks: treating cells with Nocodazole (Noco) to depolymerize microtubules and Latrunculin A (LatA) to disrupt actin filaments (Fig. 2a) [23]. Angular distribution analysis revealed strikingly different responses to the two treatments: LatA treatment left the angular distribution largely unchanged, with peaks near 0° and 180° preserved, indicating that actin filaments are dispensable for super-diffusive events (Fig. 2b, top left panel). In stark contrast, Noco treatment led to a modest reduction in the peak near 0° and a corresponding increase in the peak near 180°, suggesting decreased directional motion and increased confinement (Fig. 2b, top right panel). Most dramatically, quantitative analysis of super-diffusive event frequencies revealed that LatA treatment had no impact on the fraction of condensates exhibiting super-diffusion, while Noco treatment completely abolished super-diffusive events (Fig. 2b, bottom panels). The effectiveness of Noco in disrupting microtubules was confirmed through SiR-tubulin staining after 30 minutes of treatment [24], which demonstrated extensive microtubule depolymerization (Fig. 2c). These findings establish that super-diffusion of HOPS condensates is absolutely dependent on microtubules but independent of actin filaments, strongly implicating microtubule-based motor proteins as the driving mechanism for active transport.

**Fig. 2.**
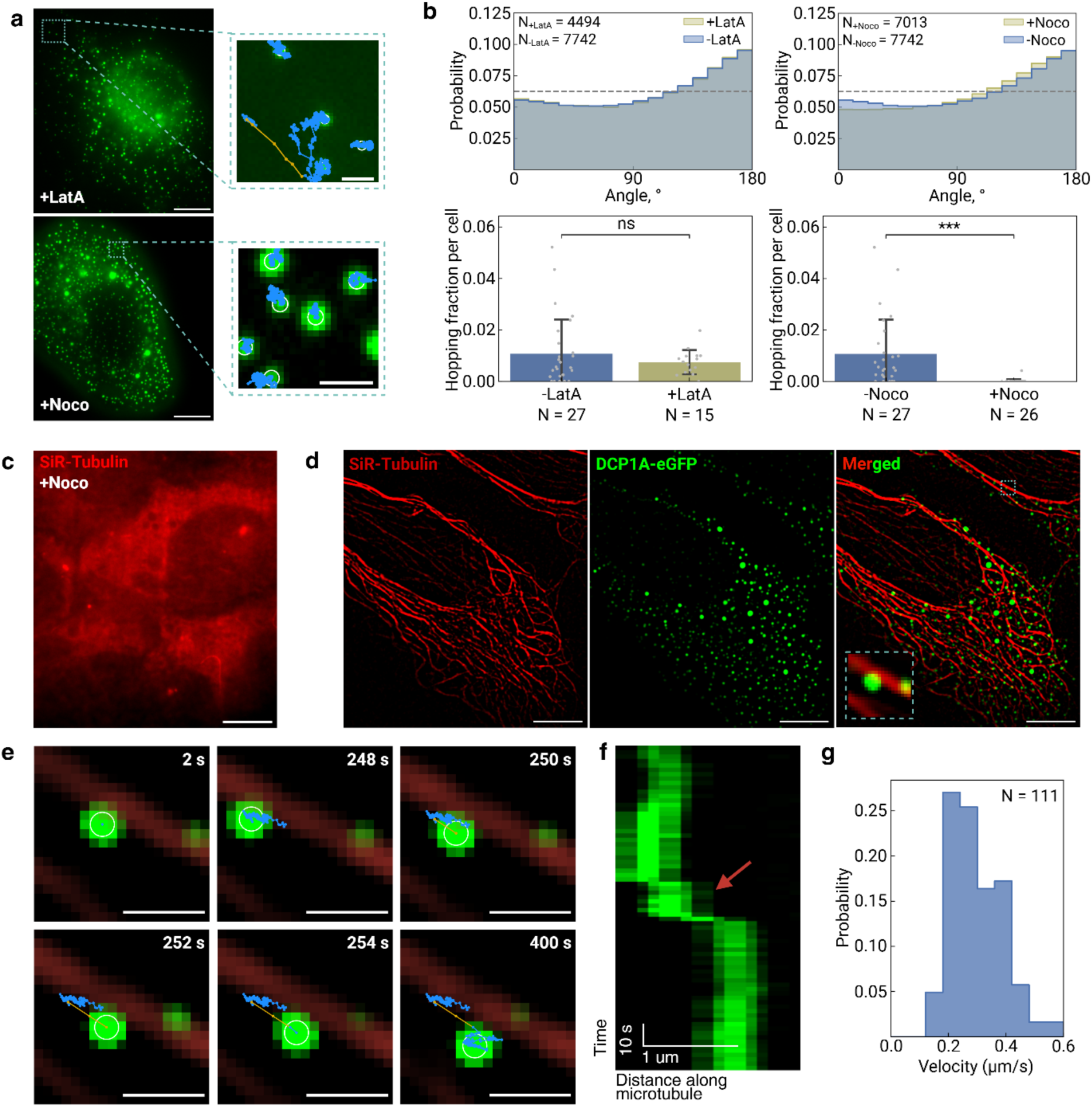
Super diffusion of HOPS condensates relies on microtubule but not actin filaments. **a |** Representative images of eGFP-labeled DCP1A in hyperosmotically compressed U2OS cells treated with latrunculin A (LatA) or nocodazole (Noco), with a zoomed-in view to show DCP1A HOPS condensates dynamics over an extended imaging window of 6.5 min. Trajectories of HOPS condensates are labeled as blue for sub diffusion while gold for super diffusion. Scale bars are 10 µm. Same below unless otherwise noted. **b |** Diffusion metrics extracted from HOPS condensate trajectories, including the distribution of angles between steps within a trajectory and a quantiXication of fraction of HOPS condensate trajectories that contains the super diffusion behavior. **c |** Representative image of a U2OS cell stained for microtubules using SiR-Tubulin under Noco treatment, showing a complete disassembly of microtubules. **d |** Dual-color imaging of SiR-Tubulin-stained microtubules and eGFP-labeled DCP1A HOPS condensates in the same cell. **e |** Time-lapse zoomed-in images of a representative HOPS condensate trajectory highlighted in **d**. Scale bar is 1 µm. **f |** Kymograph of **e** showing temporal dynamics of the super diffusion step. **g |** Velocity distribution of HOPS condensate steps during super diffusion.

The selective dependence on microtubules prompted us to directly test whether super-diffusive condensates move along microtubule tracks through live-cell co-imaging experiments. Using dual-color fluorescence microscopy, we simultaneously visualized DCP1A HOPS condensates and microtubules to assess spatial and temporal correlations during super-diffusive events (Fig. 2d). Direct observation revealed that during super-diffusive episodes, HOPS condensates moved in close proximity to and along the direction of nearby microtubules, providing compelling visual evidence for microtubule-guided transport (Fig. 2e). Quantitative analysis of a representative super-diffusive trajectory demonstrated the characteristic signature of motor protein transport: the condensate remained stationary from 0 to 248 seconds with a velocity of 0.013 μm/s, underwent rapid translocation between 248-252 seconds at 0.16 μm/s (highlighted by the red arrow in the corresponding kymograph, Fig. 2f), and then stabilized again at 0.023 μm/s for the remainder of the observation period. This velocity profile—alternating between slow, confined motion and rapid, directional transport—is characteristic of cargo transported by motor proteins along cytoskeletal tracks. Population-level analysis of all super-diffusive events revealed a narrow velocity distribution centered around 0.2 μm/s (Fig. 2h), which closely matches the velocities reported for kinesin and dynein motor proteins transporting various cellular cargoes along microtubules. Together, these direct visualization experiments provide definitive mechanistic evidence that the rare super-diffusive events represent genuine microtubule-based active transport, enabling HOPS condensates to rapidly relocate between cellular regions where they are constrained.

### HOPS condensates are not confined by physical corral

Having established the microtubule-dependent mechanism underlying super-diffusion, we next investigated the origin of the predominantly sub-diffusive behavior exhibited by HOPS condensates. One leading hypothesis for this sub-diffusion was physical corralling—the idea that cellular structures such as membranes create closed compartments that physically confine condensate movement, potentially enhanced by cytoplasmic reorganization following cell volume compression under hyperosmotic stress. To rigorously test whether such physical barriers restrict DCP1A HOPS condensate mobility, we employed 40-nm neutral Genetically Encoded Multimeric (GEM) particles as molecular probes to map spatial accessibility in hyperosmotically compressed cells [25]. GEMs are self-assembling homomultimeric scaffolds fused to sapphire that form bright particles of defined dimensions without interacting with subcellular structures, making them ideal controls for detecting physical barriers. The logic was straightforward: if DCP1A HOPS condensates are confined by membrane-bound corrals that create physically closed spaces, then GEM particles—being unable to cross such barriers—should outline these corrals by avoiding the regions they enclose and exhibit similar restricted mobility patterns.

Remarkably, we observed a striking dichotomy in mobility patterns: GEMs exhibited unrestricted diffusion throughout the cytoplasm, while DCP1A HOPS condensates remained spatially constrained to specific regions (Fig. 3a). Reconstruction of the cytoplasmic landscape using particle trajectories revealed that GEM particles could access nearly the entire cytoplasmic volume, whereas HOPS condensates were restricted to localized areas without being excluded by physical barriers (Fig. 3b-d). Quantitative analysis demonstrated this dramatic difference: GEM particles were entirely mobile, while DCP1A HOPS condensates were almost entirely immobile under identical conditions (Fig. 3e). Most strikingly, comparison of theoretical and experimental diffusion coefficients showed that GEM particles exhibited close agreement between predicted and observed values, while DCP1A HOPS condensates displayed over 100-fold lower diffusion coefficients than predicted, even for condensates with radii similar to GEMs (Supplementary Table 1). These findings definitively rule out physical corralling and establish that DCP1A HOPS condensates undergo constrained—not confined—diffusion, driven by specific molecular interactions rather than physical entrapment. This distinction is critical: constrained diffusion results from specific binding or tethering interactions that restrict movement while maintaining spatial accessibility, whereas confined diffusion involves physical barriers that create closed compartments. Having eliminated physical barriers as the cause of restricted mobility, we next aimed to identify the specific sub-cellular interactions responsible for constraining HOPS condensate movement within the accessible cytosolic space.

**Fig. 3.**
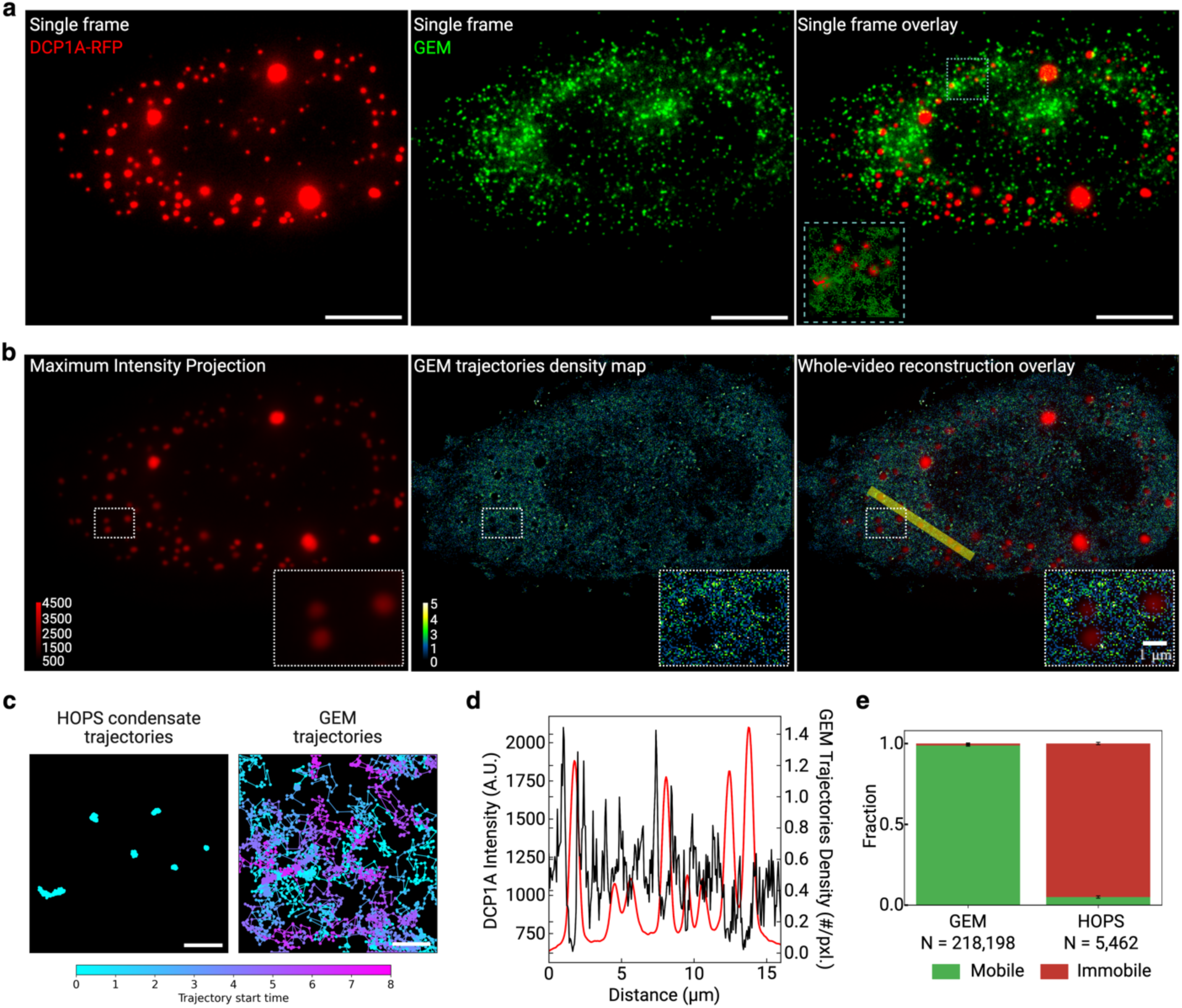
Reconstruction of GEM tracking reveals that HOPS condensate confinement is not caused by physical corrals in live cell. **a |** Representative single-frame images of a U2OS cell expressing RFP-labeled DCP1A and Sapphire-labeled GEM particles. The zoomed in view shows all the individual localizations of HOPS condensates (red) and GEM particles (green) throughout the whole duration of the movie. Scale bars are 10 µm. **b |** Reconstructed images of the same cell in **a**, with the more static HOPS condensates represented by maximum intensity projection (MIP) over time and the more dynamic GEM particles represented by a density map of the average localization of each trajectory. The density map reconstruction was performed on a grid size of 58.5 nm. **c |** Trajectories in the zoomed in region in **a**, where the trajectories are color coded based on the start time of the trajectories, showing that GEM trajectories reconstruction is from multiple particles that come into the focus plane at different times. **d |** A cross-section of DCP1A MIP intensity and GEM trajectory density per pixel over distance across the yellow line shown in **b**. **d |** QuantiXication of the mobile versus immobile fractions of HOPS condensates and GEM particles.

### Sub diffusion of HOPS condensates is caused by ER attachment

Having definitively ruled out physical corralling as the mechanism underlying HOPS condensate confinement, we next aimed to identify the specific molecular interactions responsible for their constrained diffusion through systematic screening of subcellular organelles. To broadly survey membrane associations without the bias of fluorescent labeling, we employed holotomography imaging—a non-invasive, label-free technique that visualizes intracellular structures based on their refractive index differences with the surrounding environment [26]. Since cytoplasmic signals in holotomography predominantly correspond to membrane structures, including the endoplasmic reticulum and mitochondria, this approach enabled comprehensive mapping of condensate-membrane relationships. Strikingly, DCP1A HOPS condensates demonstrated strong spatial association with membrane structures, with nearly no condensates present in membrane-devoid regions, suggesting that membrane interactions might be important to their subcellular organization (Fig. 4a). Given that HOPS condensates appeared uniformly distributed throughout the cytoplasm in close proximity to membranes, and considering that the ER forms the most extensive membrane network spanning the entire cytoplasmic volume and has been shown to regulate condensate dynamics before[27, 28], we hypothesized that ER attachment represents the primary mechanism underlying condensate confinement. This hypothesis was further supported by prior evidence that processing bodies exhibit dynamic contacts with ER membranes in U2OS cells, suggesting an evolutionary precedent for DCP1A-ER interactions that could be amplified under osmotic stress conditions.

**Fig. 4.**
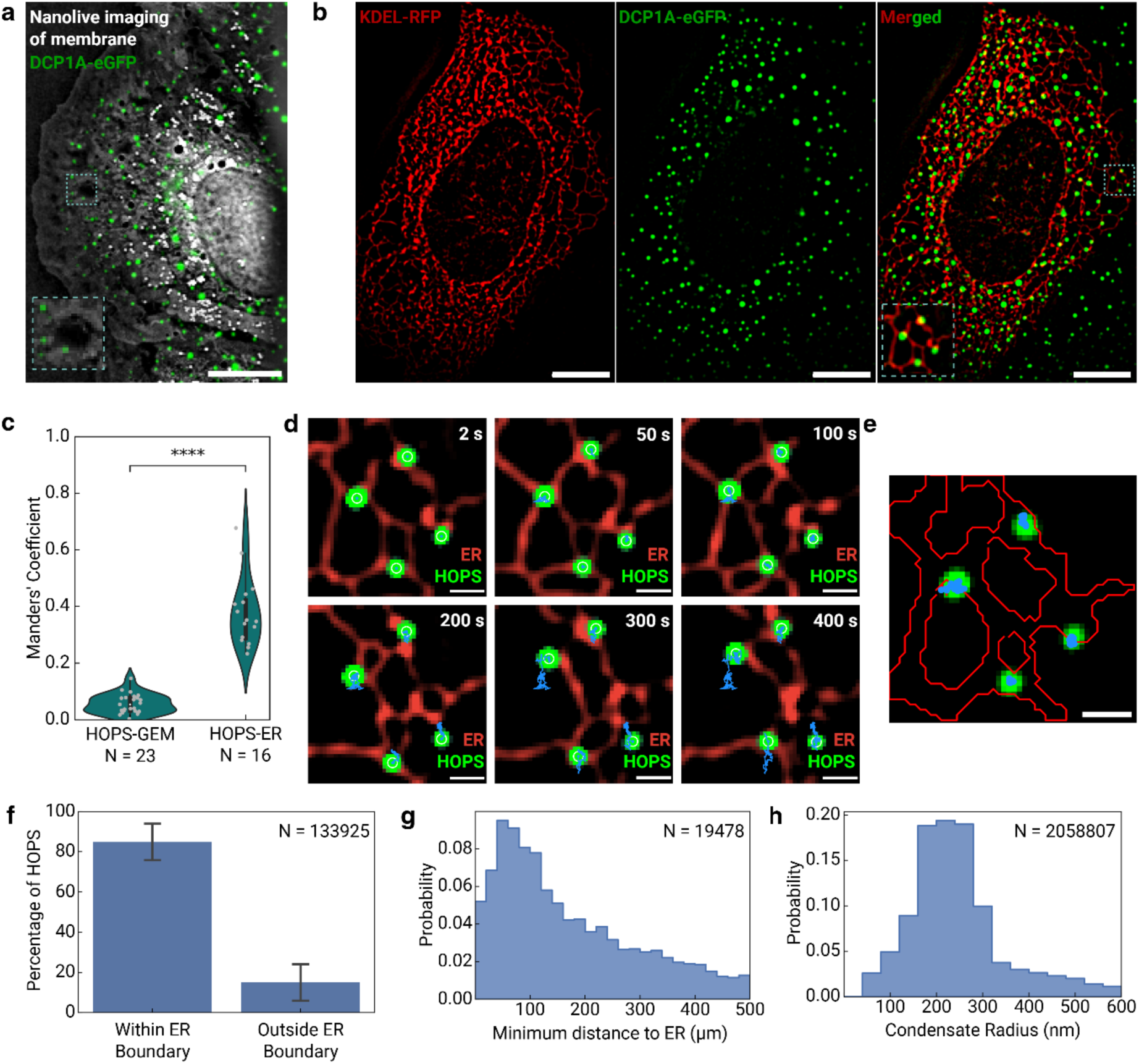
Sub diffusion of HOPS condensates is caused by ER attachment. **a |** Representative image of a U2OS cell expressing eGFP-labeled DCP1A imaged with Xluorescence microscopy and holotomography at the same time, with a zoomed-in view highlights the colocalization of HOPS condensates (green) with membrane structures that show high refractive index signal (gray). Scale bar is 10 µm. **b |** Representative image of a U2OS cell co-expressing eGFP-labeled DCP1A and ER marker KDEL-RFP. Scale bars are 10 µm. **c |** QuantiXication of colocalization between HOPS condensates with ER using Manders’ coefXicient and GEM particles as a control. **d |** Zoomed-in time-lapse images of a sub-cellular region highlighted in **b**, showing HOPS condensate dynamics relative to the ER. Scale bars are 1 µm. **e |** ER membrane boundaries segmented using ilastik overlaid with HOPS condensate trajectories from the same region. Trajectories are limited to 50 seconds to minimize photobleaching. Scale bar is 1 µm. **f |** QuantiXication of the percentage of HOPS condensates located inside versus outside the ER membrane. **g |** Distribution of distances from the center of each HOPS condensate to the nearest ER membrane boundary. **h |** Radius distribution of individual HOPS condensates.

To directly test the ER attachment hypothesis, we implemented a dual-labeling strategy by co-transfecting UGD cells with RFP-tagged ER markers (Calreticulin-KDEL) alongside endogenous eGFP-DCP1A [24], enabling simultaneous visualization of ER membranes and HOPS condensates under hyperosmotic conditions. Live-cell imaging revealed that the vast majority of DCP1A HOPS condensates localize in immediate proximity to ER membranes, with condensates appearing to track closely with the complex, branched ER network throughout the cytoplasm (Fig. 4b). Quantitative analysis using Manders’ coefficients—which measure fluorescence channel overlap [29]—demonstrated significant spatial association between HOPS condensates and ER membranes (mean coefficient ≈ 0.4), while colocalization with inert GEM particles yielded negligible overlap values as expected for a negative control (Fig. 4c). This spatial proximity is maintained dynamically over extended periods, with condensates remaining associated with ER membranes even as both structures undergo morphological changes (Fig. 4d). These findings provide compelling evidence that ER attachment represents a specific, biologically relevant interaction rather than random spatial correlation, establishing the foundation for understanding the mechanistic basis of condensate confinement.

To achieve precise quantitative measurements of the ER-condensate relationship, we employed machine learning-based segmentation using Ilastik to detect ER membrane boundaries with high accuracy [30, 31], followed by detailed analysis of condensate positioning relative to these boundaries (Fig. 4e). Trajectory analysis over 50-second intervals—optimized to minimize photobleaching artifacts—revealed that approximately 80% of HOPS condensates have their centers located within ER membrane boundaries, while the remaining 20% are positioned immediately outside (Fig. 4f). Remarkably, distance measurements for condensates with centers outside ER boundaries showed a narrow distribution peaking at ∼100 nm from the nearest ER membrane (Fig. 4g). Given that the average condensate radius is approximately 220 nm (Fig. 4h), even those condensates with centers positioned outside ER boundaries maintain direct physical contact with ER membranes, indicating universal ER tethering across the condensate population. These quantitative measurements definitively establish ER attachment as the molecular mechanism underlying the constrained diffusion of HOPS condensates, resolving the mechanistic basis for their restricted mobility in hyperosmotically compressed cells.

## Discussion

Our systematic investigation of HOPS condensate dynamics reveals a previously uncharacterized organizational principle that fundamentally challenges the prevailing view of condensates as static entities under osmotic stress. Through extended single-particle tracking, we demonstrate that HOPS condensates exhibit a sophisticated dual-mode behavior: predominant ER-tethered constraint punctuated by rare but highly directional microtubule-dependent transport events (Fig. 5). This discovery establishes that the cytosolic environment of hyperosmotically compressed cells remains dynamic and accessible rather than corralled, with condensate organization achieved through specific molecular interactions rather than physical entrapment. The biological significance of these findings extends far beyond biophysics to fundamental aspects of cellular adaptation, gene expression regulation, and kidney physiology [6], positioning HOPS dynamics as a critical component of the rapid stress response machinery that facilitates cellular adaptation under osmotic challenge.

**Fig. 5.**
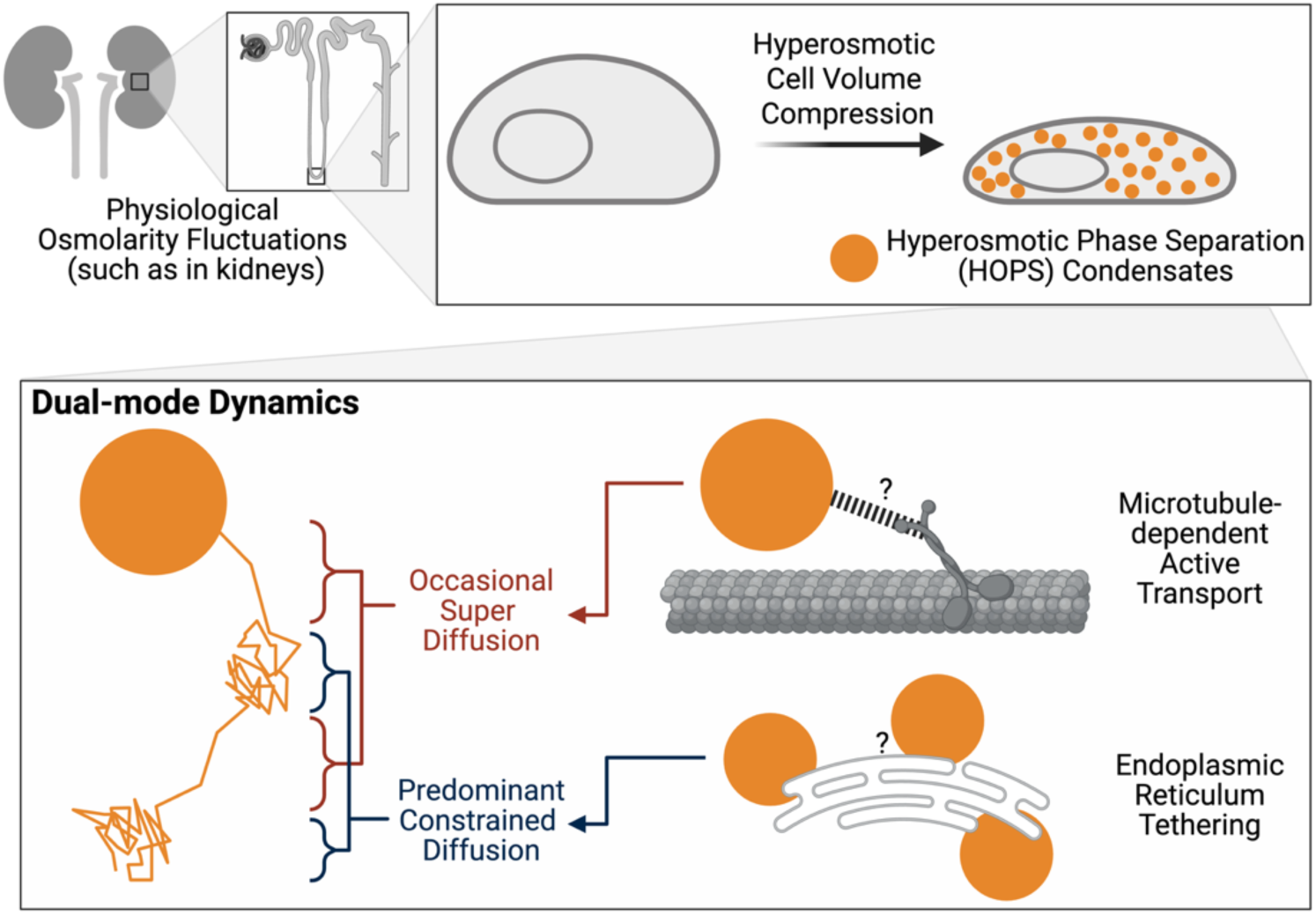
Dual-mode dynamics of HOPS condensates in hyperosmotically compressed cells.

The physiological relevance of these findings is particularly striking in the context of kidney function, where cells routinely experience dramatic osmolarity fluctuations ranging from 300 mOsm/L in the cortex to over 1,200 mOsm/L in the inner medulla [7]. Our demonstration that HOPS condensates can be rapidly repositioned via microtubule-dependent transport while maintaining ER attachment provides a mechanism for spatial reorganization of the stress response machinery in response to the steep osmotic gradients characteristic of renal physiology. The dual-mode dynamics we observe—with ∼2% of condensates undergoing active transport at velocities consistent with motor protein function—suggests a system capable of both local constraint and rapid redistribution as cellular demands change. This organizational flexibility may be essential for kidney cells to maintain function across the dramatic osmotic landscapes they inhabit, providing a physical basis for the cellular resilience required in the challenging environment of the renal medulla.

Our methodological advances have broader implications for understanding condensate behavior across biological systems. The demonstration that extended imaging duration, rather than high temporal resolution, is critical for capturing the full spectrum of condensate dynamics challenges current experimental approaches and reveals previously hidden behaviors that occur on longer timescales. The GEM particle mapping strategy we developed provides a powerful tool for distinguishing between confined and constrained diffusion, enabling researchers to identify the underlying mechanisms responsible for condensate organization in diverse cellular contexts. These approaches should prove valuable for investigating how other membraneless organelles— including nuclear condensates involved in transcription, splicing, and chromatin organization— integrate with cellular architecture to execute their biological functions.

Our identification of ER tethering as the constraint mechanism provides another instance of how different condensate-organelle interfaces serve distinct cellular functions across biological contexts [28, 32, 33]. This contrasts with recent findings in neurons, where RNA granules employ ANXA11 as a molecular tether to “hitchhike” on actively transported lysosomes, enabling long-distance mRNA delivery to distal axonal sites where local protein synthesis is required [34]. While neuronal systems prioritize condensate mobility to distribute mRNAs across long distances from soma to synapses, our hyperosmotic cell compression model system demonstrates a fundamentally different strategy: ER tethering constrains HOPS condensates rather than enabling long-range transport. This distinction reflects the different cellular challenges these systems address—neurons require efficient long-range RNA transport for spatial distribution, while cells experiencing osmotic stress employ localized constraint mechanisms that maintain condensates in specific cellular regions without the need for long-distance transport.

## Conclusions

In conclusion, we have discovered that HOPS condensates exhibit dual-mode dynamics under hyperosmotic stress—predominant constrained diffusion caused by ER tethering and rare (∼2%) super-diffusive transport events mediated by microtubule-dependent active transport. Quantitative analysis revealed that 80% of condensates have centers within ER boundaries while the remainder are positioned within 100 nm of ER membranes, establishing universal ER attachment across the population. In contrast, super-diffusive condensates move along microtubule tracks at velocities (∼0.2 μm/s) consistent with motor protein transport. Through GEM particle mapping, we demonstrated that the cytosolic space of hyperosmotically compressed cells remains highly accessible rather than corralled, challenging prevailing assumptions about the physical environment during osmotic stress. How general these dual-mode dynamics apply to other HOPS-forming proteins beyond DCP1A and whether similar ER-tethering mechanisms operate in primary kidney cells will be important questions for future studies. Our methodological advances—particularly extended imaging duration and GEM accessibility mapping—provide new tools for investigating condensate-organelle interactions across diverse cellular contexts. We thus suggest that dual-mode condensate dynamics should now be considered broadly in other models of cellular stress response organization.

## Methods

### Cell Culture and Transfections

U2OS (HTB-96, ATCC) and U2OS-eGFP-DCP1A (UGD) cell lines were cultured in McCoy’s 5A medium (Thermo Fisher, #16600082) supplemented with 10% (v/v) fetal bovine serum (Fisher Scientific, #MT35016CV) and 20 U/ml Penicillin-Streptomycin (Invitrogen, #15140122) at 37°C in 5% CO2. UGD cells were maintained under continuous G418 selection (100 μg/mL) to ensure eGFP-DCP1A construct retention. For co-imaging experiments, U2OS cells were co-transfected with GEM and DCP1A-RFP plasmids using GenJet™ In Vitro DNA Transfection Reagent (Ver. II, SignaGen Laboratories, SL100489). Briefly, 3 μL GenJet reagent was diluted in 50 μL serum-free DMEM and mixed with 0.5 μg/μL of each plasmid, incubated for 15 minutes, then added dropwise to cells at 70-80% confluency. Cells were incubated for 48 hours post-transfection before imaging.

### Cytoskeleton Disruption and Fluorescent Labeling

For cytoskeletal disruption experiments, cells at 70-80% confluency were treated with either 20 μM Nocodazole (Cayman Chemical, 31430-18-9) or 5 μM Latrunculin A (Cayman Chemical, 76343-93-6) for 30 minutes prior to imaging, with drugs maintained at identical concentrations during imaging. Microtubules were visualized using 500 nM SiR-tubulin dye (CY-SC002) added to culture medium for 4 hours at 37°C. ER membranes were labeled using CellLight™ ER-RFP BacMam 2.0 reagent (Thermo Fisher Scientific), with 100 μL added to cells at 50-60% confluency and incubated for 20-24 hours. For all imaging experiments, cells were transferred to phenol red-free Leibovitz’s L-15 medium supplemented with 10% FBS, with osmotic conditions adjusted using isotonic (8 mL L-15, 0.2 mL 10× PBS, 1.8 mL DI water) or hypertonic (8 mL L-15, 1.2 mL 10× PBS, 0.8 mL DI water) formulations.

### Fluorescence Microscopy

Live-cell imaging was performed using a Nanoimager (Oxford) equipped with a 100× objective lens, dual-band (498-524 nm and 550-620 nm) and single-band (665-705 nm) emission filters, and a 640 nm long-pass dichroic mirror channel splitter. An sCMOS camera provided 117 nm pixel resolution. Highly inclined and laminated optical (HILO) microscopy was employed at 52° excitation angle to enhance signal-to-noise ratio. Standard HOPS condensate imaging was conducted at 37°C using 0.5 Hz acquisition frequency, 100 ms exposure time, and 473 nm laser excitation at 8% power (∼2 mW) for 200 frames. Co-imaging experiments employed Alternative Light Excitation (ALEX) protocols: microtubule-HOPS co-imaging used 405 nm (8% power) and 640 nm (2% power) lasers with 500 ms exposures; GEM-HOPS co-imaging used 473 nm (10% power) and 532 nm (8% power) lasers at 50 Hz with 20 ms exposures; ER-HOPS co-imaging used 405 nm (8% power) and 532 nm (3% power) lasers with 500 ms exposures. Combined holotomography and fluorescence imaging was performed using the Nanolive 3D Cell Explorer-fluo with a 60× objective, FITC filter, 6 Hz acquisition frequency, media refractive index of 1.333, and 100 ms fluorescence exposure time.

### Holotomography and Combined Imaging

Combined holotomography and fluorescence imaging was performed using the Nanolive 3D Cell Explorer-fluo equipped with a 60× objective lens, FITC filter, and an incubator for live cell imaging. Cells were plated on a glass bottom 35 mm dish and incubated overnight. The following day, cell media was replaced with the imaging media, and cells were transferred to the microscope for imaging. The imaging frequency for combined holotomography and fluorescence was 6 Hz. The media refractive index of 1.333 was used for holotomography calculations, and an exposure time of 100 ms was used for fluorescence imaging. This label-free technique visualizes intracellular structures based on their refractive index differences with the surrounding environment, with cytoplasmic signals predominantly corresponding to membrane structures including the endoplasmic reticulum and mitochondria.

### Single-Particle Tracking and Trajectory Extraction

Video preprocessing involved band-pass filtering using difference of Gaussian (DoG) filters with sigma values optimized for HOPS condensates (1 and 5 pixels) and processing bodies (3 and 7 pixels) [31]. Spot detection and trajectory extraction were performed using TrackMate with Laplacian of Gaussian (LoG) detection (5-pixel estimated diameter, 585 nm), Linear Assignment Problem (LAP) tracking algorithm, and maximum linking distance of 5 pixels (585 nm). The spot quality threshold was set based on the peak of false-positive spots in the quality value distribution. Trajectories were exported as CSV files and analyzed using custom Python scripts for diffusion profiling [35, 36]. To determine immobile particles, the mean step size of each trajectory was calculated and compared to a threshold of 30 nm, based on a static localization error of 16 nm determined using immobile mRNA molecules attached to glass surfaces. Trajectories with mean step sizes below 30 nm were excluded from further analysis.

### Running Window Diffusion Analysis

To distinguish between confined and unconfined mobile particles, anomalous diffusion exponents (α), diffusion coefficients (D), and step sizes were computed using running window analysis. Each trajectory was divided into overlapping segments of 20 frames, with parameters calculated for each segment. The anomalous diffusion exponent (α) was determined by fitting the mean squared displacement (MSD) versus lag time (τ) curve on a log-log scale using:

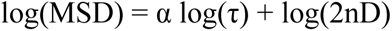

where n represents the number of dimensions (n = 2) and D is the diffusion coefficient. The optimal number of fitting points for MSD-τ analysis was approximately half the trajectory length. Trajectories with poor fits (R² < 0.6) were excluded from calculations. The diffusion coefficient (D) for each window was obtained by fitting the MSD curve to: MSD = 2nDτα. Mean step sizes were computed for each window, and angles between consecutive steps were calculated with values ranging from 0° (no directional change) to 180° (complete directional reversal).

### Super-Diffusive Event Detection

Super-diffusive steps were identified using combined criteria: threshold step size of 2 pixels (234 nm) for two consecutive steps, total displacement threshold of 5 pixels (585 nm), α threshold > 1, diffusion coefficient (D) threshold > 0.01 μm²/s, and R² > 0.6. Trajectory segments meeting these criteria were flagged as super-diffusive events for subsequent analysis.

### Co-Imaging Analysis and Channel Separation

For DCP1A HOPS and microtubule co-imaging, channel alignment was performed using bead maps collected on the same day, converted into matrices for alignment correction. Following alignment, HOPS and microtubule movies were separated for individual analysis. For DCP1A HOPS and GEM co-imaging, both signals were collected alternately in the green channel, separated by subtracting consecutive frames since DCP1A HOPS and GEM particles were cross-excited by 473 nm laser while only DCP1A HOPS was visible with 532 nm laser. Trajectories were plotted separately and analyzed to determine mobile and immobile fractions. For DCP1A HOPS and ER co-imaging, signals were acquired alternately in the green channel with DCP1A HOPS and ER appearing in separate, alternating frames.

### Colocalization Quantification

Manders’ coefficients were calculated using ImageJ plugins to quantify spatial association between HOPS condensates and ER membranes. These coefficients measure fluorescence channel overlap, with values approaching 1 indicating high colocalization and values near 0 indicating minimal overlap. GEM particles served as negative controls for colocalization analysis.

### Boundary Detection and Segmentation

ER membrane and condensate boundaries were segmented using Ilastik [30], a machine learning-based pixel classifier, given it’s been profiled to be the optimal boundary detection method for condensates with various sizes [31]. For each experimental condition, training datasets of at least 15 cells were used, with pixels classified into condensate and background classes using human visual assessment as ground truth. The trained model was applied to predict segmentation masks using Ilastik’s “simple segmentation” mode in export settings. The findContours function was employed to extract detected boundaries from segmentation masks. This ML-based approach provides robust and accurate boundary detection for cellular structures, particularly condensates, which can be challenging to detect using traditional image processing techniques.

### Spatial Analysis and Distance Measurements

Trajectory analysis was performed over 50-second intervals to minimize photobleaching artifacts. Distance measurements for condensates positioned outside ER boundaries were calculated to the nearest ER membrane. Condensate radii were measured from individual particle analysis. Spatial reconstructions were generated using trajectory density mapping with 58.5 nm grid resolution. The viscosity of cytoplasmic space under isotonic and hypertonic conditions was estimated using the Stokes-Einstein equation based on GEM particle diffusion coefficient distributions, which were then used to calculate theoretical diffusion coefficients for comparison with experimental values.

## Acknowledgements

We sincerely thank Qi (Archie) Geng and Kristen Verhey for their valuable advice on the design of the cytoskeleton-perturbation experiments. We are also grateful to Alexander Johnson-Buck and Adrien Chauvier for their insightful discussions throughout the project development and paper writing. Additionally, we thank the Biointerfaces Institute for providing access to the Nanolive 3D Cell Explorer.

## Declarations

### Consent for publication

Not applicable.

