## Supplementary Information for "ER tethering and active transport govern condensate diffusion during hyperosmotic stress"

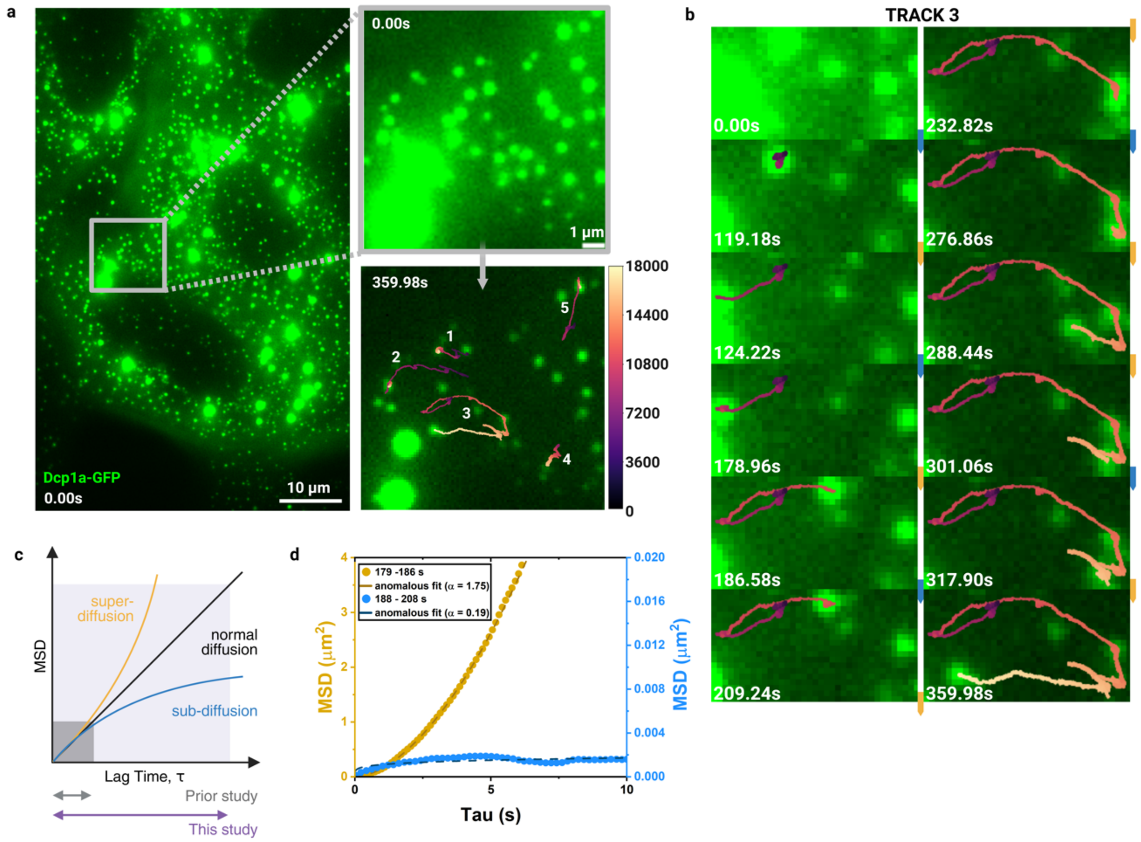


**Fig. S1 | Dual-mode dynamics of hyperosmotic phase separation (HOPS) condensates reproduced under 50 Hz imaging frequency**

**a |** Representative images of U2OS cells expressing eGFP-labeled DCP1A under hyperosmotic stress conditions. Enlarged views emphasize trajectories of hyperosmotic stress-induced HOPS condensates, where individual particle locations throughout a trajectory are colored by frame number. **b |** Time-lapse zoomed-in images of a representative HOPS condensate trajectory (#3 shown in **a**), showing both predominant sub diffusion and the occasional super diffusion. **c |** The mean squared displacement (MSD) as a function of lag time (τ) demonstrates the ability of extended time scales to differentiate between various diffusion modes. **d |** Comparison of MSD-τ curves between the sub-diffusive portions and the super diffusion portions of the same set of HOPS condensate trajectories.

**Table S1 | Theoretical and experimentally determined diffusion coefficient of HOPS condensate and GEM particles from four representative particles each**

| | Radius (nm) | Theoretical D ($\mu m^2/s$) | Experimental D ($\mu m^2/s$) |
|---|---|---|---|
| HOPS Particle #1 | 80.84 | 0.0394 | 0.000232 |
| HOPS Particle #2 | 186.70 | 0.0170 | 0.000189 |
| HOPS Particle #3 | 272.16 | 0.0117 | 0.000319 |
| HOPS Particle #4 | 313.11 | 0.0101 | 0.000118 |
| GEM Particle #1 | 40 | 0.0797 | 0.136530 |
| GEM Particle #2 | 40 | 0.0797 | 0.017967 |
| GEM Particle #3 | 40 | 0.0797 | 0.022489 |
| GEM Particle #4 | 40 | 0.0797 | 0.014519 |
